# Farmyard Mulching in Regenerative Agriculture Enhances Saprotrophs and Concomitantly Reduces Pathogenic Fungal Genera

**DOI:** 10.1101/2024.08.14.607896

**Authors:** Pratyusha Naresh, Indira Singh

## Abstract

Regenerative agriculture (RA) using nature-friendly methods nurtures soil microbial communities. Indian RA farmers use diverse practices to manage their soil. This study compared the fungal communities in RA plots with those in conventional agriculture (CA) and barren land (BL) plots (comprising completely barren-BL and with Eucalyptus - BL-Euc). Two crops were considered - Finger millets and Vegetables (Tomato/ beans) for this study. ITS metagenomic analysis of soil DNA samples obtained from RA, CA and BL plots was done to identify fungal composition in each of the study plots. The fungal communities in RA finger millet and RA vegetable were compared with respective CA finger millet and CA vegetable and with BL plots. Vegetable RA plots observed higher abundances of fungal Operational Taxonomic Units (OTUs) than in CA vegetable and BL plots. Whereas the RA finger millet plots had similar fungal OTUs as in CA finger millet and BL plots. The vegetable RA plot carrying out natural farming for 12 years (maximum length in our samples) recorded the highest fungal OTU (13707) levels while the CA plots had average OTU abundance of (7416). RA plots in both crops showed a significant reduction in plant pathogenic fungal genuses - *Bipolaris* and *Pyrenochaetopsis*. Furthermore, RA finger millet plots showed an enhanced representation of saprotrophs while CA (finger millets) had pathotroph-saprotrophs suggesting a favorable increase in decomposer populations in RA.

## Introduction

With the world moving towards sustainability in agriculture through natural/ regenerative practices, soil and plant microbial health is gaining more and more prominence. Healthy soil boasts of millions of different species of bacteria, fungi, earthworms, nematodes, insects and other invertebrates (De Gannes et al., 2016; Costantini and Mocali, 2022; Bach et al., 2020). These organisms together help to build the soil’s texture, porosity, water retention capacity, and quality through nutrient cycling processes, in effect supporting plant growth and productivity (Schröder et al., 2016).

Fungi play significant roles in dictating soil health and agricultural outcomes (Frąc et al., 2023). In an agricultural ecosystem, fungi carry out essential processes involving decomposing the organic matter and enriching the soil with minerals, making them accessible by the plant roots by solubilizing these minerals. In addition, they release several chemicals such as siderophores and indole acetic acid promote plant growth, and associate with plant roots enabling better mineral and water uptake, and finally also protect plants against pests and pathogens (Selvasekaran and Chidambaram, 2020). Many fungi are directly associated with the plants as endophytes enhancing the plant’s health and productivity (Poveda et al., 2022). Several others get into symbiotic relationships with plants and algae such as lichens, which help plants by supporting soil formation, nitrogen fixation, and fixing carbon dioxide through the photosynthesizing algae (Du et al., 2019). However, there is also a host of pathogenic fungi that end up destroying the crops and reducing their productivity (Doehlemann et al., 2017). Interestingly, nature provides us with a plethora of diverse solutions to overcome pests and pathogenic species. Among the fungal kingdom also a large number of species have been recorded with anti-pathogenic properties (Pandit et al., 2022). Climate change has been overburdening the pest problem (Hyde, 2022). Hence, in recent scenarios analysis of the soil fungi is essential to arrive at nature-based solutions for resolving the climate mediated agricultural challenges.

Regenerative agriculture uses a wide variety of soil enriching practices that dissuade the use of chemicals but rely on organic inputs such as farmyard waste, compost, crop mulch etc. These dead organic matters form a rich resource for the growth and multiplication of saprophytic microbes. Going with this hypothesis this research assessed the fungal community structure in natural/ regenerative agriculture (RA) and compared it with the conventional agricultural plots and barren land and eucalyptus containing plots. Like the bacterial populations in RA soils which were seen to host a more diverse community of bacteria with increase in prominent sets of Plant Growth Promoting Rhizobacteria (PGPR) genera, similarly here we have explored the impacts of healthier agriculture practices on fungal profiles.

## Materials and Methods

### Soil Sampling and Chemical Analysis

Two crops were considered for the study: ragi (finger millet) and vegetables (tomato/beans). This study is an extension of the previously published - Singh et. al., 2023, which analyzed the bacterial community structures in the plots. We have three types of plots - Regenerative (RA), Conventional (CA) and Barren Land (BL). The details of the soil samples, the region’s soil type and the history of the land have been provided in Table 1. Here we have analyzed the fungal profiles in these plots. The soil chemical profiling for each of the samples was carried out as discussed in Singh et. al., 2023. The soil chemical profiles of the respective soil samples are provided in Table 2.

**Table 1.** Soil sampling, classification, agricultural practices and history.

| Type of Crop | Sample Names | Place of Soil Sample Collection with (Latitude/ Longitude/ Altitude) in m | Soil Classification (as per USDA soil taxonomy) | Type of Agriculture | Previous Crop | No of Plots | No of Years of Practice | Major Regenerative Agriculture Practices | History of the plot |
| --- | --- | --- | --- | --- | --- | --- | --- | --- | --- |
|  | Con-VP1 and Con-VP2 | Ramanagara –<br>Con-VP1 - 12.785677/ 77.218218/ 747 m<br>Con-VP2 - 12.785252/ | Fine, Mixed, Isohyperthermic, Kandic Paleustalfs | Conventional | Con-VP1 - Crop rotation with leafy vegetables. Con-VP2 – | 2 | - | Use of NPK fertilizers and chemical pesticides along with | Both Con-VP1 and ConVP2 have been following mixed |
| Vegetable<br>(Beans/<br>Tomato<br>) |  | 77.217067/<br>747 m |  |  | crop<br>rotation<br>with<br>carrots |  |  | farmyard<br>manure | agricult<br>ure<br>practice<br>by<br>using<br>both<br>farmyar<br>d<br>manure<br>and<br>chemica<br>l<br>treatme<br>nt to<br>support<br>soil and<br>plant<br>health<br>for 5-10<br>years.<br>The<br>adjacen<br>t plots<br>use<br>regener<br>ative<br>practice<br>s and<br>intercro<br>pping<br>and<br>agrofor<br>estry<br>etc<br>which<br>would<br>also<br>impact<br>these<br>plots. |
|  | Reg-1VA | Magadi<br>12.956508/<br>77.190900/<br>925 m | Clayey-skeletal,<br>Mixed,<br>Isohyperthe<br>rmic,<br>Kandic<br>Paleustalfs | Regene<br>rative | None as<br>the<br>land<br>was<br>fallow<br>since<br>last ten<br>years.<br>This | 1 | 1 | Cow<br>dung,<br>vermi-<br>compost,<br>and<br>Jeevamru<br>tha, crop<br>rotation<br>and | The<br>plot<br>was a<br>Paddy<br>and<br>Ragi<br>grown<br>land a<br>long |
|  |  |  |  |  | was the first crop |  |  | inter-cropping; Seed treatment with Beejamrutha. Since this was the first crop they did not do crop residue management, slight till of 3-6 inches was done. | time ago but no farming was being done on this plot since 2018. The land was acquired by the current farmer in 2020. He prepared it for doing agriculture and followed the practices as given in adjacent column. This was the first crop that this farmer had grown. |
|  | Reg-3VP | Magadi 13.031895/77.326871/ | Clayey-skeletal, Mixed, Isohyperthermic, Kandic | Regenerative | Green leafy vegetables | 1 | 3 | Farm manure and Jeevamrutha applied | Farm plot was uncultivated for 20 |
|  |  | 925 m | Paleustalfs |  |  |  |  | twice a year and mixed-cropping and crop rotation, Neem oil for pest control. Seed treatment with *Pseudomonas* and *Trichoderma* | years. For the past 3 years regenerative agriculture was being practiced as given in adjacent column. Farmer has been growing vegetables and fruit trees from past five years |
|  | Reg-8VP | Ramanagara<br>12.783899/<br>77.216949/<br>747 m | Fine, Mixed, Isohyperthermic, Kandic Paleustalfs | Regenerative | Chilli | 1 | 8 | Farm manure, and Jeevamrutha applied twice a year, mixed cropping, crop rotation and Beejamrutha 3-6 inch minimal tillage and using rotovator | This farmer has been consistently practicing regenerative agriculture as discussed in adjacent column for 8 years. |
|  |  |  |  |  |  |  |  | for managin<br>g crop<br>residue<br>and then<br>it is<br>added<br>with<br>farm<br>manure<br>followed<br>by<br>mulching | Before<br>that for<br>12<br>years<br>he was<br>doing<br>inter-<br>croppin<br>g<br>growin<br>g caster,<br>red<br>gram,<br>hyacint<br>h<br>beans,<br>cowpea<br>and<br>sorghu<br>m. |
|  | Reg-10VP | Hosur<br>12.609869/<br>77.702348/<br>880 m | Loamy,<br>Mixed,<br>Isohyperthe<br>rmic,<br>Typic<br>Rhodustalfs | Regene<br>rative | Horse<br>Gram | 1 | 10 | 400 kg<br>Farm<br>manure<br>per bed<br>twice a<br>year and<br>Jeevamru<br>tha<br>through<br>drip and<br>spray,<br>mulching<br>and<br>Panchgav<br>ya. Crop<br>rotation<br>with<br>legumes.<br>For some<br>seeds<br>*Pseudom*<br>*onas*<br>treatmen<br>t was<br>given* | This<br>large<br>farm<br>was<br>previou<br>sly used<br>for<br>cultivati<br>on of<br>vegetab<br>le crops<br>for<br>3 years<br>by<br>natural<br>method<br>s. For<br>the<br>past ten<br>years<br>since<br>the land<br>was<br>acquire<br>d by the<br>current<br>owners,<br>regener<br>ative agriculture has been practiced here as given in adjacent column. |
|  | Reg-12VA | Ramanagara<br>12.785475/<br>77.219237/<br>747 m | Fine, Mixed, Isohypermic, Kandic Paleustalfs | Regenerative | Sesbania bispinosa | 1 | 10 -12 | 4-5 tons Farm manure and Vermicompost, per year, Jeevamrutha and microbial culture added monthly twice during crop growth; multi-cropping with crop rotation; seed treated with Beejamrutha and cow urine. Crop residue is managed with rotovator and eventually used as green manure. Minimal till of 3-6 | Previously the land was conventional agriculture land. For past 12 years the farmer adopted regenerative agriculture. He has been growing multiple varieties of ragi, rice etc. He has developed a seed-bank. Now he grows multiple |
|  |  |  |  |  |  |  |  | inches. | varieties of vegetables and greens and fruits using regenerative agriculture methods as outlined in adjacent column |
| Ragi | Con-RA1 & Con-RA2 | Doddaballapur<br>Con-RA1-13.377086/77.543199/880 m<br>Con-RA2-13.374351/77.549439/880 m | Fine, Mixed, Isohyperthermic, Typical Haplustepts | Conventional | Con-RA1 - The plot was a mango plantation before the current ragi crop<br>Con-RA2 - Ragi with mixed crop as red gram, cowpea, castor, sorghum, hyacinth beans etc. | 2 | - | Use of NPK fertilizers and chemical pesticides alone. No other supplementation | The plots have been doing ragi cultivation for the last 3 – 5 years using conventional methods. Another disadvantage for these plots is their vicinity to eucalyptus plantations in their neighboring plots. |
|  | Reg-1RA | Magadi<br>12.998967/<br>77.306946/<br>925 m | Clayey-skeletal, Mixed, Isohyperthermic, Kandic Paleustalfs | Regenerative | Ragi with mixed crop as red gram, cowpea, castor, sorghum and hyacinth beans etc are grown year after year | 1 | 1 | Farm manure and green manure, mulching ; natural insecticide for pest management. This was first year no crop residue or mulching done. Minimal till of 3-6 inches. | It was an abandoned land for a long time until the current farmer acquired the land and started the cultivation using regenerative agriculture methods as given in adjacent column |
|  | Reg-7RA | Ramanagara<br>12.779501/<br>77.269773/<br>747 m | Fine, Mixed, Isohyperthermic, Kandic Paleustalfs | Regenerative | Horse gram | 1 | 7 | cow dung, jaggery, Vermicompost, Jeevamrutha; A special organic pesticide + cow urine spray for pest management; crop rotation with legumes, crop rotation; seed treatment with Beejamrutha, cow dung, jaggery and calcium for seed treatment; organic pest management | Initially the farmer grew mango and ragi the conventional way. Later he also tried to grow mulberry but that failed. For the past seven years he has been practicing regenerative agriculture as detailed in adjacent column |
|  | Reg-8RA | Ramanagara<br>12.783881/<br>77.217184/<br>747 m | Fine, Mixed, Isohyperthermic, Kandic Paleustalfs | Regenerative | Ragi with Mixed crop such as red gram, cowpea, castor, sorghum and hyacinth beans etc. grown year after year | 1 | 8 | Farm manure, green leaves manure and Jeevamrutha applied twice a year; crop rotation with leguminous crops; seed treatment with cow urine; pest management also with cow urine and natural<br>pesticide | This farmer has been consistently practicing regenerative agriculture as discussed in adjacent column for 8 years. Before that for 12 years he was doing inter-<br>croppin<br>g<br>growin<br>g castor,<br>red<br>gram,<br>hyacint<br>h<br>beans,<br>cowpea<br>and<br>sorghu<br>m. |
| Barren<br>Land | *BL-Euc*<br>*& BL* | Doddaballapur<br>13.376992/<br>77.542568/<br>880 m | Fine, Mixed,<br>Isohyperthermic,<br>Typic<br>Haplustepts | Not<br>applica<br>ble | - | 2 | - | No<br>treatmen<br>t | The<br>land<br>has<br>either<br>been<br>lying<br>barren<br>for 20<br>years or<br>more or<br>planted<br>with<br>eucalyp<br>tus |

Note: In the provided names the following nomenclature has been followed –
*Reg* - Regenerative; *Con* - Conventional; *BL* - Barren Land. *V* - Vegetable; *R* - Ragi; *P* - soil sampling in Presence of crop; and *A* - soil sampling in Absence of the crop; and the numbers after the hyphen indicate the number of years of Regenerative agriculture practice.
Jeevamrutha – composed of soil, chickpea flour, jaggery, cow dung and cow urine; Panchagavya – composed of milk, butter, curd, cow dung and cow urine; Beejamrutha – comprises of cow dung, cow urine, soil and lemon juice.
"Reprinted with permission from 'Regenerative agriculture augments bacterial community structure for a healthier soil and agriculture' by Singh et al. (2023), licensed under CC-BY, https://www.frontiersin.org/articles/10.3389/fagro.2023.1134514/full"

**Table 2:** Chemical Parameters of the Soil Samples.

| Sample Name | pH | EC (dS/m) | Organic carbon (%) | Nitrogen | Phosphorus | Potassium | Calcium | Magnesium | Zinc | Manganese | Iron | Copper |
| --- | --- | --- | --- | --- | --- | --- | --- | --- | --- | --- | --- | --- |
|  |  |  |  | kg/ha |  |  | mEq/1000 g |  | ppm |  |  |  |
| **IDEAL** | **6.5 - 7.5** | **<1.00** | **0.5-0.75** | **280-560** | **22.9 - 56.33** | **141-336** | **>1.5** | **>1.0** | **>0.6** | **>2.0** | **2.5 - 4.5** | **>0.2** |
| Con-VP1 | 7.54 | 0.367 | 0.39 | 131.8 | 342.47 | 250.3 | 38 | 21 | 3.72 | 7.2 | 35.64 | 1.17 |
| Con-VP2 | 7.6 | 0.399 | 0.44 | 106.6 | 346.28 | 284.4 | 41 | 27 | 4.77 | 7.68 | 11.31 | 0.84 |
| Reg-1VA | 7.41 | 0.113 | 0.29 | 125.4 | 87.64 | 217.9 | 35 | 19 | 1.23 | 6.33 | 13.32 | 0.42 |
| Reg-3VP | 7.43 | 0.193 | 0.3 | 156.8 | 62.39 | 184 | 40 | 30 | 4.44 | 3.99 | 32.16 | 1.56 |
| Reg-8VP | 8.31 | 0.267 | 0.36 | 120.1 | 152.8 | 189.7 | 62 | 47 | 2.34 | 6.27 | 8.79 | 0.72 |
| Reg-10VP | 7.95 | 0.279 | 0.51 | 144.2 | 510.13 | 506.4 | 105 | 64 | 4.08 | 8.34 | 19.62 | 2.19 |
| Reg-12VA | 7.71 | 0.231 | 0.32 | 100.3 | 187.33 | 223.5 | 79 | 54 | 2.79 | 9.42 | 16.53 | 0.81 |
| Con-RA1 | 5.79 | 0.316 | 0.35 | 131 | 37.6 | 334 | 14 | 6 | 1.32 | 10.8 | 16.56 | 0.45 |
| Con-RA2 | 3.94 | 0.159 | 0.39 | 144 | 44.7 | 170 | 21 | 10 | 1.14 | 18.63 | 78.27 | 1.5 |
| Reg-1RA | 7.35 | 0.128 | 0.38 | 106.6 | 58.58 | 216.6 | 58 | 32 | 1.05 | 11.01 | 10.32 | 0.51 |
| Reg-7RA | 6.89 | 0.09 | 0.36 | 119.1 | 148.61 | 252.7 | 45 | 28 | 2.58 | 14.58 | 36.18 | 1.56 |
| Reg-8RA | 7.01 | 0.13 | 0.42 | 125.4 | 60.01 | 306.7 | 54 | 38 | 2.37 | 16.02 | 23.13 | 0.57 |
| BL-Euc | 6.04 | 0.235 | 0.31 | 119 | 29 | 242 | 27 | 14 | 1.51 | 23.91 | 9.3 | 0.459 |
| BL | 5.85 | 0.106 | 0.41 | 150 | 14.7 | 108 | 22 | 9 | 0.99 | 6.33 | 16.29 | 0.327 |

Note: The ideal values are based on recommendations given by the Indian Society of Soil Science (31). "Reprinted with permission from 'Regenerative agriculture augments bacterial community structure for a healthier soil and agriculture' by Singh et al. (2023), licensed under CC- BY, https://www.frontiersin.org/articles/10.3389/fagro.2023.1134514/full"

### DNA isolation, Quality check and Sequencing

Soil DNA isolation was performed using the DNeasy Power soil kit, following the manufacturer’s protocol. Quality check for the isolated DNA was done using Nanodrop, by estimating A_260_/A_280_ ratio. These steps are discussed in detail in Singh et. al., 2023. This was followed by internal transcriber spacing (ITS) based sequencing, for fungal metagenomic analysis. Sequencing was done using Illumina MiSeq platform (Eurofins Genomics India Pvt. Ltd., Bangalore, India). ITS rRNA gene amplification was done using the following primers: ITS 1F (5’-GCATCGATGAAGAACGCAGC-3’) and ITS 2R (5’-TCCTCCGCTTATTGATATGC-3’).

### Metagenomic Analysis and Statistics

The obtained sequences were subjected to metagenomics analysis using QIIME 2 pipeline (Bolyen et al., 2019). This pipeline uses the DADA2 algorithm for the crucial de-noising step, which aids in filtering and trimming reads that do not meet the quality requirements. This is followed by identification of amplicon sequence variants (ASVs). These ASVs are then annotated using a trained classifier model to assign taxonomy. Following this, a thorough literature review was conducted to analyze and understand the results obtained and identify relevant taxa. Statistical analysis was performed on these using unpaired t-tests and one-way ANOVA using Python Programming Language.

### Functional analysis: FUNGuild

FUNGuild v1.1 was used to investigate the function of fungal communities in soil samples (Nguyen et. al., 2016). Twelve functional guilds, namely, ectomycorrhizal fungi, fungal endophytes, wood saprotrophs, plant saprotrophs, dung saprotrophs, undefined saprotrophs, animal pathogens, plant pathogens, algal parasites, animal parasites, fungal parasites and lichen parasites were identified. These were classified according to three trophic modes (pathotrophs, saprotrophs and symbiotrophs). Three confidence ranks, “possible”, “probable”, and “highly probable”, were assigned by the tool. One-way ANOVA was performed using python programming to assess the effect of regenerative agriculture on the relative abundance of OTUs inferred using FUNGuild.

## Results

### Metagenomics

ITS metagenomics studies were carried out for all 14 soil samples given in Table 1. The sequence information for the samples can be found in NCBI with the project ID PRJNA912401 (accession numbers SRR28254635 to SRR28254648). A total of 4,656,196 raw sequence reads were obtained from the metagenomic library generated using Illumina. This ranged from 257,444 to 438,222 reads per sample. These were processed to remove primers and other noise and chimeras, to obtain 3,382,950 high quality reads. Fungal taxonomy up to the genus level have been considered for this study. Taxonomic analysis revealed 3907 OTUs in the samples.

### Abundance of Fungal Populations

In vegetable RA plots we found a relatively higher abundance of fungal OTUs (Reg-8VP: 11097, Reg-3VP: 9569, Reg-1VA: 8901, Reg-10VP: 9569 and Reg-12VA: 13707) as compared to that in CA (Con-VP1: 5597 and Con-VP2: 9235) and BL (BL-Euc: 8447 and BL: 9630) (Figure 1A). Whereas the fungal OTUs in RA finger millet plots were found to be lesser or equivalent to the CA plots (Figure 1B).

**Figure 1:**
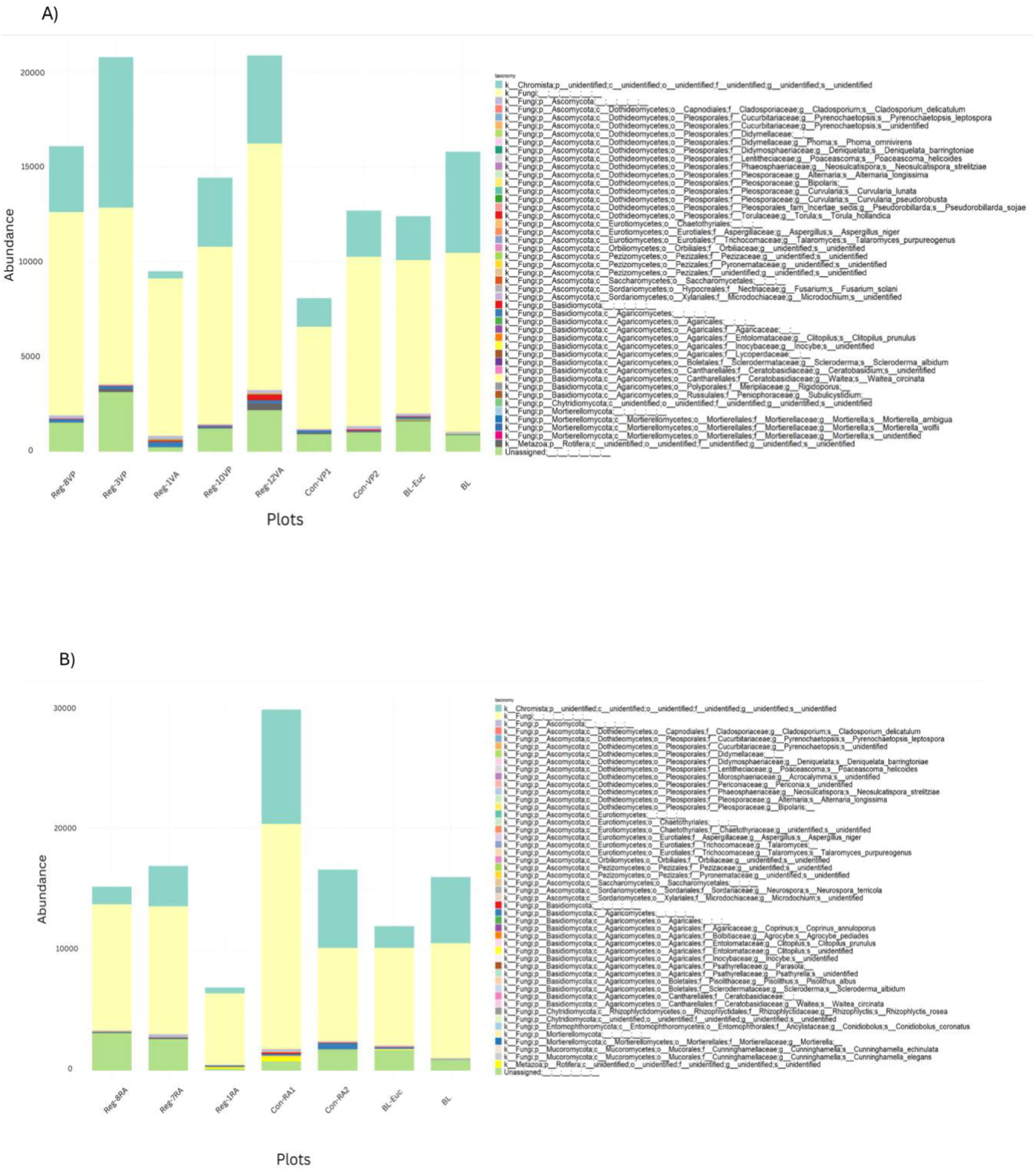
Total fungal abundance in A) Vegetable (RA, CA) and BL plots; and in B) Finger millet (RA, CA) and BL plots

### Abundance of Beneficial Fungi

The finger millet and vegetable RA plots were compared with their respective CA controls and BL negative controls for the abundance of beneficial fungi known in literature. Here we have considered the well characterized Plant Growth Promoting Fungi (PGPFs), endophytic fungi and ectomycorrhizal fungi which are well known beneficial genera. Based on the identified taxa in our samples the PGPFs considered in this study include - *Cladosporium*, *Aspergillus*, *Talaromyces*, *Inocybe*, *Psathyrella*, *Rhizophlyctis*, *Conidiobolus*, *Mortierella*, *Phoma* and *Pseudorobillarda* (Hossain & Sultana, 2020; Adedayo & Babalola, 2023; Li et al., 2024; Halifu et al., 2021; El-Katatny, 2010; Gałązka et al., 2020). Endophytes identified were *Acrocalymma*, *Agrocybe*, *Clitopilus*, *Cunninghamella*, *Torula* and *Subulicystidium* (Fite et al., 2023; Khasim et al., 2020; Atiphasaworn et al., 2017; Syamsia et al., 2021; Ma et al., 2023), while the ectomycorrhizal fungi were *Pisolithus* and *Scleroderma* (Singh et al., 2019) (Figure 2).

**Figure 2:**
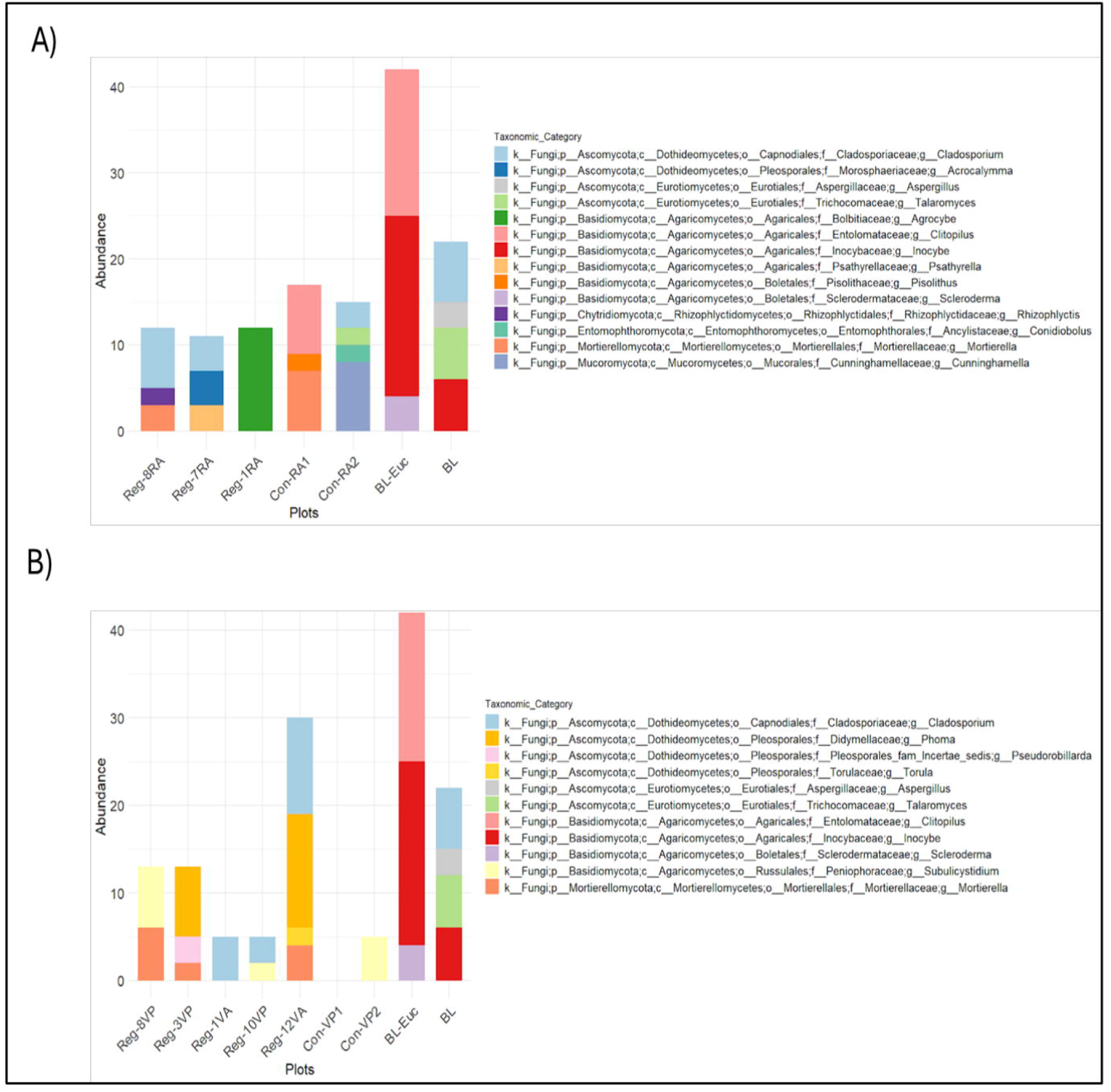
Abundance (Number of Reads) of beneficial fungi across RA, CA and BL plots with A) Comparing Finger millet plots and B) Comparing Vegetable plots

Each of the three RA finger millet plots were found to contain specific beneficial fungal genera - *Rhizophlyctis* (0.01%) in Reg-8RA, *Acrocalymma (*0.02%) in Reg-7RA and *Agrocybe* (0.18%) in Reg-1RA respectively, which were absent in the finger millet CA and BL plots (Figure 2A). In addition, Reg-8RA also contained *Cladosporium* (0.05%) and *Mortierella* (0.02%), while Reg-7RA also showed presence of *Cladosporium* (0.02%) and *Psathyrella (*0.02%). Interestingly, the BL-Euc (Eucalyptus) soil showed highest levels of beneficial fungal genera - *Scleroderma (*0.03%), *Clitopilus (*0.14%) and *Inocybe* (0.18%). Among the vegetable RA plots, the one carrying out regenerative agriculture for 12 years, Reg-12VA showed enhanced representation of beneficial fungal genera - *Mortierella* (0.02%), *Phoma* (0.06%), *Torula* (0.01%) and *Cladosporium* (0.05%) (Figure 2B).

### Abundance of Pathogenic Fungi

Simultaneously the occurrence and abundance of plant pathogenic fungal genera, chosen based on their occurrence across the samples - *Pyrenochaetopsis* (Kwaśna et al., 2021), *Deniquelata* (Ariyawansa et al., 2013, Devadatha et al., 2018), *Periconia* (Gunasekaran et al., 2021), *Alternaria* (Tozlu et al., 2018, Pinto & Patriarca, 2016), *Bipolaris* (Al-Sadi, 2021, Aslam et al., 2021, Naik et al., 2016), *Microdochium* (Gagkaeva et al., 2020), *Waitea* (Vojvodić et al., 2022, Vojvodić et al., 2021), *Curvularia* (Aguas et al., 2017, Wei et al., 2022), *Fusarium* (Rampersad, 2020, Blanco & Aveling, 2018), *Ceratobasidium* (Li et al., 2024, De Melo et al., 2018) and *Rigidoporus* (Chaiharn et al., 2019, Mahendran et al., 2021) was explored and compared in RA, CA and BL plots (Figure 3).

**Figure 3:**
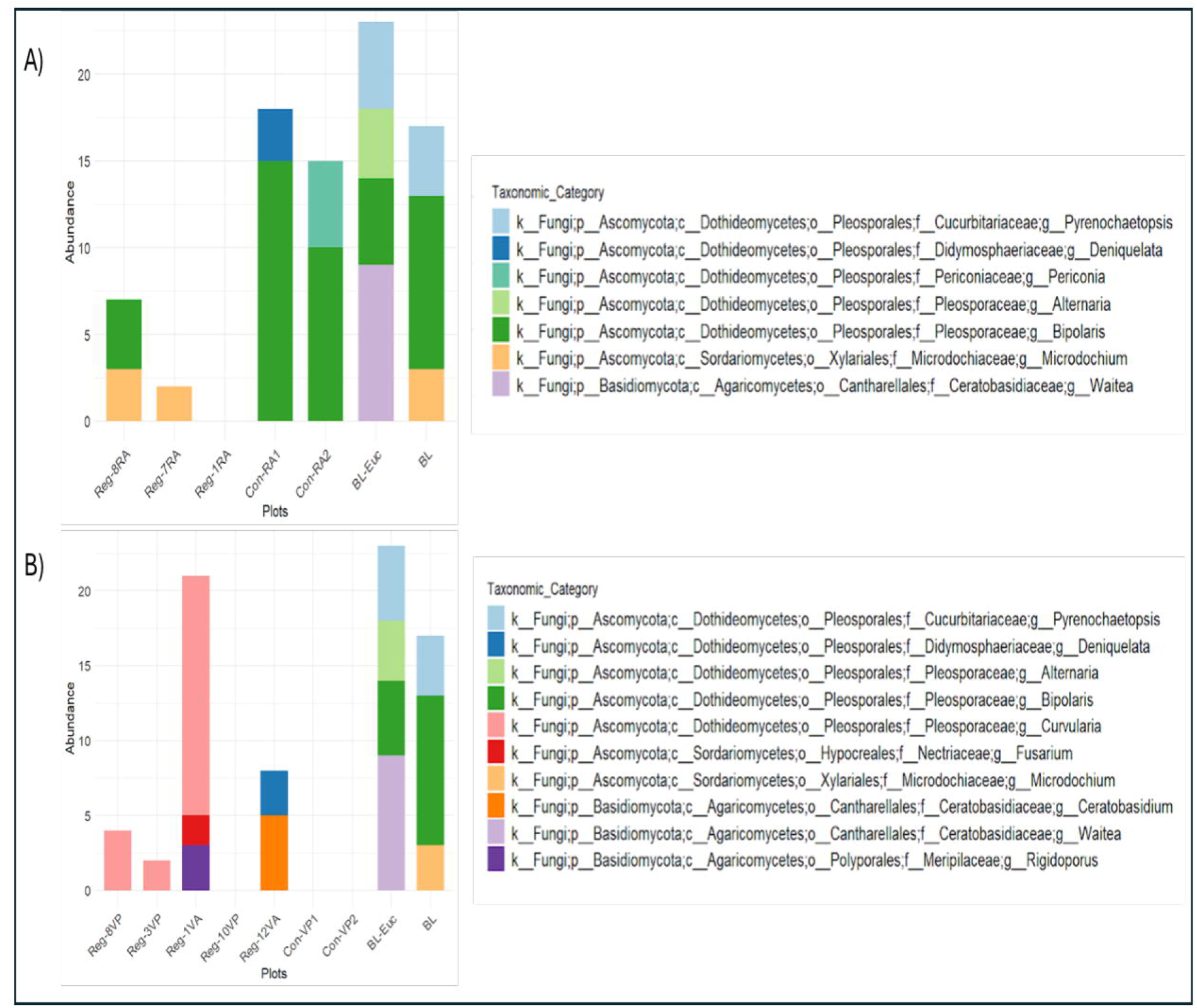
Abundance (Number of Reads) of plant pathogenic fungi across RA and CA vegetable plots and BL plots in A) Finger millet and B) Vegetable plots

The RA finger millet plots showed an absence of several of the plant pathogenic fungal genera. Reg-8RA was found to contain *Bipolaris* (0.03%) and *Microdochium* (0.02%), while Reg-7RA had only *Microdochium* (0.01%) and Reg-1RA contained none of the plant pathogenic genera. Clearly, the finger millet RA (0.05%, 0.01% and 0%) plots were found to contain fewer of the plant pathogenic fungal genera compared to finger millet CA (0.06% and 0.09%) and BL (0.19% and 0.11%) plots. The well-known plant pathogen genus *Bipolaris* in finger millet CA plots was found to be (0.05% and 0.06%) and in the BL plots (0.04% and 0.06%) (Figure 3A). Statistical analysis (one-way ANOVA) was done which revealed that there is significant decrease in the total abundance of pathogenic fungi in case of finger millet (p-value = 0.0397)

In the case of vegetables, the total pathogenic fungal genera in each RA plot (0.02%, 0.01%, 0.22%, 0% and 0.03%) were found to be mostly lesser than that in the BL plots (0.19% and 0.11%). However, the vegetable CA plots did not contain any of the pathogenic fungal genera that we analyzed for this study. Interestingly, none of the vegetable RA or CA plots had *Bipolaris* or *Pyrenochaetopsis* while the BL plots contained (0.04% and 0.06%) *Bipolaris* and (0.04% and 0.03%) *Pyrenochaetopsis* (Figure 3B). Statistical analysis revealed that abundance of the pathogenic fungal genera *Bipolaris* and *Pyrenochaetopsis*, have decreased significantly in RA plots both in case of finger millets (p = 0.0355 and p = 0.00077) and vegetables (p= 0.00195 and 0.000017) for *Bipolaris* and *Pyrenochaetopsis* respectively.

### Functional Analysis

The identified fungal taxa were classified into ecological-guilds (trophic modes) using the FUNGuild tool to understand their relative distribution patterns in the various agricultural systems versus barren plots. The taxa could be classified into five trophic modes - pathotroph-saprotroph-symbiotrophs, pathotroph-saprotrophs, saprotrophs, saprotroph-symbiotrophs and symbiotrophs. Finger millet RA samples were found to be composed of 71 % saprotrophs, 19% pathotroph-saprotrophs and 10% saprotroph-symbiotrophs, whereas the CA comprised 70% pathotroph-saprotrophs, 14% saprotrophs, 12% saprotroph-symbiotrophs and 4% symbiotrophs and BL had 58% pathotroph-saprotrophs, 38% symbiotrophs, and 4% pathotroph-saprotroph-symbiotrophs (Figure 4A). The RA plots show a higher abundance of saprotrophs and lesser levels of pathotroph-saprotrophs while both CA and BL plots are observed to have greater relative abundance of the latter.

**Figure 4:**
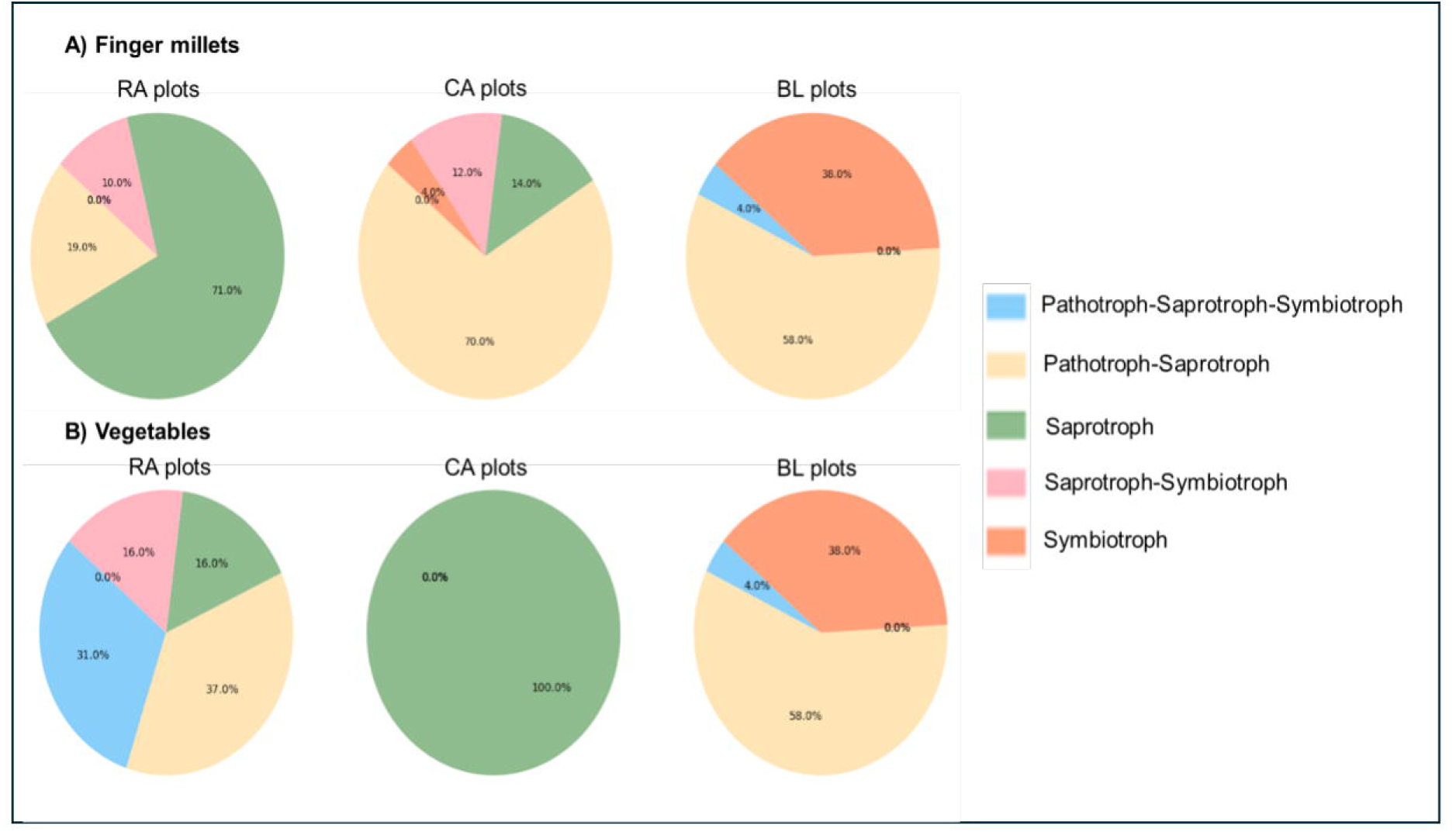
Relative abundance of trophic modes in Regenerative agriculture (RA), Conventional agriculture (CA) and Barren land (BL) plots in A) Finger millet and B) Vegetable

In vegetables, the RA samples had 37% pathotroph-saprotrophs, 31% pathotroph-saprotroph-symbiotrophs, 16% saprotrophs and 16% saprotroph-symbiotrophs, while the CA was found to comprise of 100% saprotrophs and BL contained 58% pathotroph-saprotrophs, 38% symbiotrophs, and 4% pathotroph-saprotroph-symbiotrophs (Figure 4B). In vegetable RA plots although all five trophic modes were present the significant change observed was the additional assemblage of saprotroph-symbiotrophs which is absent in both vegetable CA and BL plots.

Statistical analysis of relative abundance of each trophic mode between the samples in A and B (Figure 4) at 95% level of significance revealed the following. In the case of finger millet plots, p-values of 0.340, 0.055, 0.477, 0.487 and 0.128 were obtained for the trophic modes Pathotroph-Saprotroph-Symbiotroph, Pathotroph-Saprotroph, Saprotroph, Saprotroph-Symbiotroph and Symbiotroph respectively. In the case of vegetable plots, p-values were 0.402, 0.002, 0.577, 0.314 and 0.034 for the trophic modes Pathotroph-Saprotroph-Symbiotroph, Pathotroph-Saprotroph, Saprotroph, Saprotroph-Symbiotroph and Symbiotroph respectively. Thus, in vegetable plots (RA and CA) versus BL plots significant differences were observed in case of pathotroph-saprotrophs and symbiotrophs. Although these results largely indicate that there is not much significant difference in the trophic modes assembly among RA, CA and BL, this could be attributed to the smaller sample size used in this study.

Aside from the trophic modes, FUNGuild was also used to assign functional groups or guilds to the OTUs. This revealed that among finger millet RA samples, most fungi belonged to saprotrophic guilds (plant saprotrophs, wood saprotrophs, undefined saprotrophs and dung saprotrophs) along with others like endophytes, algal parasites and plant pathogens. Plant pathogens were absent in Reg-7RA and Reg-1RA and comprised a lesser fraction in Reg-8RA (17.39%). Contrary to this the CA plots apart from hosting saprotrophs (undefined saprotrophs and plant saprotrophs), plant pathogens and endophytes, also contained animal parasites and animal pathogens (Figure 5A). More importantly, Reg-7RA showed higher abundance (50%) of wood saprotrophs.

**Figure 5:**
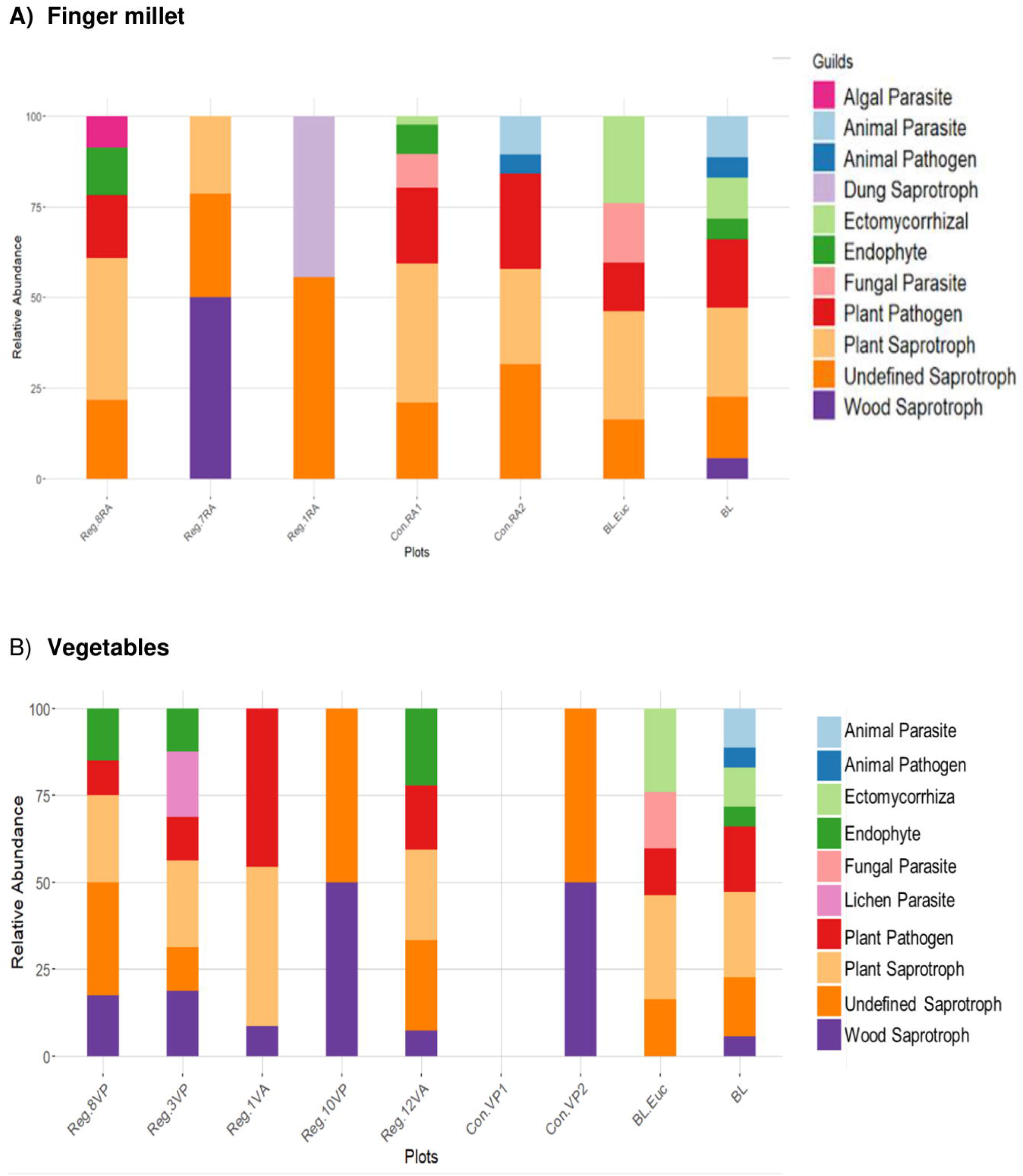
Relative abundance of functional guilds in respective RA, CA and BL plots A) Finger millets and B) Vegetable plots

Among vegetables, a large share of the RA samples consisted of saprotrophic fungi (wood, undefined and plant saprotrophs). We also observed an enhanced representation of endophytes in RA vegetable plots. In addition, RA vegetable plots also contain plant pathogens and lichen parasites. In contrast, CA (vegetable) and BL had saprotrophs, plant pathogens, fungal parasites, animal parasites, animal pathogens and ectomycorrhizal fungi (Figure 5B).

Regenerative agriculture samples in both cases have a significant percentage of saprotrophic guilds, including plant saprotrophs.

## Discussion

This is the first ever report from India that has attempted to study the soil fungal populations in regenerative agriculture versus conventional agriculture, and barren and eucalyptus-containing land. Like several other studies on fungal community profiling, we also did not find a consistent pattern of fungal assemblages with agricultural practices (Hannula, et.al., 2021; Sommermann, et.al., 2018; Schneider, et.al., 2010). Also, with many fungal species still not identified, the characterization of fungal community structures is extremely difficult. Moreover, with almost negligible information available on fungi in Indian soils, it is essential to analyze and qualify soil health and ecosystems based on fungal populations in India.

The vegetable RA plots interestingly showed higher fungal abundance compared to the vegetable CA plots. This may be due to the healthier soil management practices and organic supplementations used in RA contributing to a thriving fungal flora. In addition, the use of pesticides and herbicides in conventional agriculture has been suggested to cause unintended harmful effects on soil microorganisms (Steiner, et.al., 2024). Furthermore, the vegetable RA plot Reg-12VA, (Table 1) observed the highest fungal population. Reg-12VA also contained the highest levels of beneficial fungi among all the RA plots. These could be attributed to 12 years of natural farming practiced here with heavy input of green farm mulch which retains moisture and forms a rich carbon resource supporting the growth of decomposers especially fungi (Iqbal, et. al., 2020).

Studies also show that mulching and compost amendments reduce proliferation of pathogenic fungal strains (Entry, et. al., 2005; Neher, et. al., 2022; Vida, et. al., 2020; Richardville, et. al., 2022; Blaya, et. al., 2016). Mulch accumulates microbes with cellulose degrading properties which also act on cell walls of fungal pathogens. Conforming with these findings we also report that both vegetable and finger millet RA plots practicing for at least 3 years and more had reduced numbers of pathogenic fungal genera. Both the finger millet and vegetable RA plots showed significantly lower levels of well-known plant pathogens *Bipolaris* and *Pyrenochaetopsis*. In vegetable CA plots a complete absence of all the assessed pathogenic genera was observed, suggesting either that the usage of farmyard manure promotes the growth of pathogen suppressing strains or the use of pesticides and herbicides by these farms has contributed to their elimination.

Functional analysis of the fungal communities revealed an interesting, enhanced occurrence of saprotrophs in RA finger millet plots while CA finger millet and BL plots showed highest representation of pathotroph-saprotrophs. Literature shows that soil amendments with rich sources of organic matter stimulate proliferation of saprotrophic fungi (Clocchiatti, et. al., 2021). Regenerative agriculture practices using farmyard manure, vermicompost, cow dung, leaves and green mulch as used by the plots Reg-1RA, Reg-7RA and Reg-8RA provide the necessary organic matter that enhances saprotrophic fungal growth. Our findings suggest that the proliferating saprotrophs enrich the soil for nutrients in plant assimilable forms by decomposing organic matter. Additionally, their overactive cellulase enzymes non-specifically also digest the pathotroph-saprotrophs leading to their simultaneous decline (Nam, et. al., 2023). Thus, organic input agriculture provides us with the desired dual advantage of enhancing soil nutrients while concomitantly reducing pathogenic strains.

In contrast, the vegetable RA plots showed higher fractions of pathotroph-saprotrophs and pathotroph-saprotroph-symbiotrophs while the CA plot had the desired 100% saprotroph composition. Here we construe that since the vegetable CA plots are using organic supplementation alongside the pesticide and herbicides, the latter are damaging the fungal populations, but the organic supplements are allowing for the revival of the saprotrophs. On the contrary, in the vegetable RA plots the fungal populations are in a dynamically evolving phase which keeps changing with crops and organic supplementations and may take several years to reach a mature and more organized structure. To understand this, it is necessary to study well evolved farms that have been carrying out regenerative practice for at least 20 - 25 years.

Somehow, none of the assayed farms showed the presence of Arbuscular Mycorrhizal Fungi (AMFs) which are known to be significant supporters of plant growth and productivity. AMFs also form an important subset of the saprotrophic fungal guild. Studies suggest that among several factors that dictate enhancement of the plant’s synergistic association with AMFs are - (i) limiting nitrogen and phosphorus conditions; (ii) increased root secretion of strigolactones and flavonoids and (iii) vicinity to tree populations (Shantz, et. al., 2016; Della Mónica, et. al., 2023; Dierks, et. al., 2021). The relatively scant population of trees around most of the studied agricultural plots might have contributed to the noted absence of AMFs in our study. This stirs up the consequential debate - whether agroforestry should be the recommended practice of choice for regenerative agriculturists. Another noteworthy point was that the BL-Euc plot contained the highest levels of beneficial as well as pathogenic fungal genera. Greater association of fungi around trees is a common phenomenon, as trees provide an abundant carbon resource in the form of dead wood, dried leaves etc (Manici, et. al., 2023; Dierks, et. al., 2021).

In our previous report on bacteria in these same plots (Singh, et. al., 2023), we found that bacterial diversity had pronouncedly increased in natural agriculture farms and was directly proportional to the number of years of RA implementation. Although fungal abundance was found to increase in vegetable RA plots since we could not see a similar trend in finger millet plots, we are unable to confidently conclude this finding. This points to the need for many more experiments with larger sample sizes to verify this result. In contrast to the bacterial study where we could see a remarkable accumulation of PGPRs in RA plots, fungal study showed a decline in pathogenic fungal strains in RA plots. It is evident that fungi and bacterial populations respond differently to agricultural practices, however we can conclude that RA brings along a healthy improvement in their compositions thus enhancing the vitality of crops and their productivity.

## Author contributions

IS was involved in planning the experiment, collecting soil samples, DNA isolation, data analysis as well as writing the manuscript. PN has done the complete ITS metagenomic data analysis for the fungal community structure in plots and wrote the Materials and Methods and Results Section with the Figures in this manuscript.

## Acknowledgements

This research is part of the SHEFS - an interdisciplinary research partnership, forming part of the Wellcome Trust’s funded Our Planet, Our Health programme, with the overall objective to provide novel evidence to define future food systems policies to deliver nutritious and healthy foods in an environmentally sustainable and socially equitable manner. This research was funded by the Wellcome Trust through the Sustainable and Healthy Food Systems (SHEFS) Project (Grant number-205200/Z/16/Z). The authors would like to thank ATREE for facilitating the initial research.

